# Isoxanthohumol improves obesity and glucose metabolism via inhibiting intestinal lipid absorption with a bloom of *Akkermansia muciniphila* in mice

**DOI:** 10.1101/2023.06.21.543373

**Authors:** Yoshiyuki Watanabe, Shiho Fujisaka, Yoshitomo Morinaga, Shiro Watanabe, Allah Nawaz, Hideki Hatta, Tomonobu Kado, Ayumi Nishimura, Muhammad Bilal, Muhammad Rahil Aslam, Keiko Honda, Yoshimi Nakagawa, Samir Softic, Kenichi Hirabayashi, Takashi Nakagawa, Yoshinori Nagai, Kazuyuki Tobe

**Author notes:** Address correspondence to: Shiho Fujisaka, MD. PhD., Associate professor, First Department of Internal Medicine, Faculty of Medicine, University of Toyama, Japan Address: 2630 Sugitani, Toyama, 930-0194, Japan, Kazuyuki Tobe, MD. PhD., Professor, First Department of Internal Medicine, Faculty of Medicine, University of Toyama, Japan Address: 2630 Sugitani, Toyama, 930-0194, Japan.

## Abstract

**Aims:** Dysbiosis is an important factor that leads to metabolic disorders by disrupting energy balance and insulin sensitivity. A decrease in *Akkermansia muciniphila* is a phenotype of obesity-induced dysbiosis. Although interventions to increase *A. muciniphila* are expected to improve glucose metabolism, the underlying mechanism has not been fully understood.

**Methods:** Isoxanthohumol (IX), a prenylated flavonoid found in beer hops was administered to high fat diet-fed mice. We analyzed glucose metabolism, gene expression profiles and histology of liver, epididymal adipose tissue and colon. Lipase activity, fecal lipid profiles and plasma metabolomic analysis were assessed. Fecal 16s rRNA sequencing was obtained and selected bacterial species were used for in vitro studies. Fecal microbiota transplantation and monocolonization were conducted to antibiotic-treated or germ-free (GF) mice.

**Results:** The administration of IX lowered weight gain, decreased steatohepatitis and improved glucose metabolism. Mechanistically, IX inhibited pancreatic lipase activity and lipid absorption by decreasing the expression of the fatty acid transporter CD36 in the small intestine, which was confirmed by increased lipid excretion in feces. IX administration improved the gut barrier function and reduced metabolic endotoxemia. In contrast, the effects of IX were nullified by antibiotics. As revealed using 16S rRNA sequencing, the microbial community structure changed with a significant increase in the abundance of *A. muciniphila* in the IX-treated group. An anaerobic chamber study showed that IX selectively promoted the growth of *A. muciniphila* while exhibiting antimicrobial activity against some *Bacteroides* and *Clostridium* species. To further explore the direct effect of *A. muciniphila* on lipid and glucose metabolism, we monocolonized either *A. muciniphila* or *Bacteroides thetaiotaomicron* to GF mice. *A. muciniphila* monocolonization decreased CD36 expression in the jejunum and improved glucose metabolism, with decreased levels of multiple classes of fatty acids determined using plasma metabolomic analysis.

**Conclusion:** Our study confirmed a direct role of *A. muciniphila* in energy metabolism, which was induced by microbial actions of IX. These highlight new treatment strategies for preventing metabolic syndrome by boosting the gut microbiota with food components.

## Introduction

The gut microbiota is a core component of metabolic control [1-3]. They outnumber somatic cells in the gut and are active metabolic species, which maintain the homeostasis of biological activities by metabolizing and synthesizing lipids, amino acids, and vitamins, converting bile acids, and maintaining immune function [4]. Moreover, metabolites derived from gut microbiota act as metabolic signaling molecules. For instance, short-chain fatty acids (SCFAs) and bacterial metabolites of non-digestible dietary fiber are a source of nutrients for the intestinal epithelium and anti-obesity effects by promoting the secretion of glucagon-like peptide 1 (GLP-1) and peptide YY in intestinal L cells via G-protein-coupled receptor 43 and 41 signaling[5]. In addition, the activation of G-protein-coupled bile acid receptor (TGR5), a receptor for secondary bile acids, increases energy expenditure through thyroid hormones, promotes GLP-1 secretion, and exerts anti-inflammatory effects on macrophages[6]. Thus, the gut microbiota is inherently beneficial for energy balance, glucose metabolism, and immune system. However, under dysbiosis state caused by a high-fat diet (HFD) and obesity, the altered microbial structure disrupts metabolic regulation, leading to further obesity and insulin resistance[3]. One of the mechanisms of insulin resistance by dysbiosis is an elevated chronic inflammation derived from metabolic endotoxemia, which is driven by elevated intestinal permeability[7]. Thus, modifying microbial function to restore the energy-regulatory systems and interventions to enhance the intestinal barrier function can improve metabolic dysfunction.

Mucin-degrading *Akkermansia muciniphila*, a major species of Verrucomicrobia, resides in the mucus layer[8] in humans and rodents. The relative abundance of *A. muciniphila* decreases in obesity and type 2 diabetes[9]. *A. muciniphila* increases mucus in the intestinal epithelium and enhances the intestinal barrier to improve metabolic endotoxemia, which improves insulin resistance and prevents liver damage and atherosclerosis[10]. In addition, the oral administration of pasteurized *A. muciniphila* improves insulin sensitivity and hyperlipidemia in overweight/obese participants [11], suggesting that *A. muciniphila* is a promising microbiota as a novel therapeutic modality for metabolic dysfunction. However, an efficient way to increase the relative abundance of *A. muciniphila* and the mechanisms of host metabolic improvement are not well understood. .

Polyphenols possess antioxidant, antibacterial, and anti-inflammatory properties[12]. They also modify the microbiota to improve glucose metabolism[13, 14]. Isoxanthohumol (IX), a heat-stable prenylated flavonoid, is synthesized by isomerizing xanthohumol, which is a flavonoid unique to beer hops and is produced in the beer brewing process. IX improves HFD-induced metabolic disorders, which are associated with microbiota changes [15].

To this end, we aimed to elucidate the mechanisms of metabolic improvement by IX from the viewpoint of intestinal function and microbial actions.

## Methods

### Animal studies

Male C57BL/6 mice were purchased from Japan SLC Inc. (Tokyo, Japan). The GF C57BL/6 mice were housed in vinyl isolators and obtained by natural mating. The mice were maintained under a 12-h light-dark cycle and provided *ad libitum* access to water and food. Six-week-old mice were divided into groups and fed HFD, HFD + IX, or HFD + IX + antibiotics. The HFD was purchased from Research Diets Inc (NJ, USA). IsoxanthoFlav (IX) was obtained from Hopsteiner (Mainburg, Germany). Each diet was frozen until use and replaced weekly with a fresh diet. Energy intake was calculated by measuring the average weight of food consumed and multiplying it by the number of calories per unit weight of diet. To establish reproducibility, the experiments were conducted in several independent cohorts, and similar trends were observed. For studies involving the measurement of relatively large variations such as body weight, food intake, and glucose tolerance test results, data from several mouse cohorts were verified by considering the cage effect. In some experiments, ITT and euthanization of mice were performed at different ages, and we displayed data from representative cohorts.

In the study using GF mice, the number of mice per group varied because the mice bred in our isolator were used to ensure that the environmental and genetic backgrounds and ages among all the mice were identical. The animal care policies and experimental procedures were approved by the Animal Experiment Committee of the University of Toyama.

### Culture Experiments

Bacteria were cultured at 37 °C in an anaerobic environment (Bactron EZ, Toei Kaisha Ltd., Tokyo, Japan). Brain Heart Infusion (BHI; Becton Dickinson and Company, New Jersey, USA) was used as the culture medium. During co-culturing with IX, BHI and the bacterial solution were cultured at a ratio of 1000:1. The absorbance at 600 nm was measured after 24 h for *Escherichia coli* and after 48 h for other bacteria.

### Bacterial transplantation

*A. muciniphila* or *Bacteroides thetaiotaomicron* was cultured with BHI + four supplements (Hemin solution 0.5 mg/mL, Menadion solution 1 mg/mL, L-Cysteine solution 0.1 mg/mL, and resazurin 0.1 %) as medium at 37 °C for 48 h in an anaerobic incubator. The culture medium was centrifuged for 5 min, the supernatant was discarded, the pellet was suspended in 2 mL of PBS, and 200 µL was orally administered to mice by gavage.

### Fecal microbiota transplantation

For bacterial transfer into GF mice, fecal microbiota transplantation (FMT) was performed thrice every alternate day by gastric gavage of 200 μL filtered feces suspended in saline. Fecal samples were collected from groups of mice that were fed the HFD or HFD + IX diet for 4 weeks. All recipient mice were maintained on a normal diet after the transfer.

### Antibiotics

In the antibiotic cohort, a mixture of vancomycin (0.5 g/L), metronidazole (0.5 g/L), neomycin (0.5 g/L), and ampicillin (0.5 g/L) (Sigma-Aldrich, St. Louis, MO) was administered to mice from 6 weeks of age in drinking water.

### OGTT and ITT

Each OGTT (2 g/kg weight) and intraperitoneal ITT (1.0 unit/kg) was performed after fasting the animals for 4 h. Blood samples were collected from the tail at specific time intervals, and glucose levels were measured using a Stat Strip XP3 (Nipro, Japan).

### Bomb calorimetry

Feces were collected from each mouse for 24 h. After drying at 50 °C for 16 h, the energy contents in the feces were measured with an IKA calorimeter C6000 (IKA, Osaka, Japan) according to the manufacturer’s instructions. .

### Triglyceride and non-esterified fatty acid analyses

The feces excreted over 24 h were collected and air-dried. After the feces were weighed and pulverized with a motor and pestle, 100 mg-portions of the fecal powder were used for the extraction of total lipids according to the method of Bligh and Dyer. The fecal total lipids were applied to preparative silica gel thin–layer chromatography (TLC) plates and NEFAs and TGs were isolated by developing the plates with the mixture of petroleum ether, diethyl ether and acetic acid (80:30:1, v/v/v). Fatty acid methyl esters (FAMEs) were prepared from the NEFAs and TGs fractions, which were analyzed by gas-liquid chromatography as described previously [16].

### Analysis of TG levels in the liver and plasma

The liver and plasma TG levels were measured using a Triglyceride Colorimetric Assay Kit (Cayman Chemical Company, U.S.A.), according to the manufacturer’s instructions.

### Fecal mucin measurement

Fecal mucin levels were determined based on the fluorometric measurement of O-linked reducing sugars using a commercially available kit (Cosmo Bio, Tokyo, Japan). Images were obtained using a microscope connected to a digital camera (BZ-X800; Keyence, Osaka, Japan).

### Pancreatic lipase activity

Pancreatic lipase activity was measured based on the amount of NEFAs generated during the incubation with olive oil emulsified with 5% Gum Arabic solution and porcine pancreatic lipase. IX was dissolved at the desired concentrations in DMSO, which was added to the reaction mixture 5 min before adding porcine pancreatic lipase. The effects of the test compounds on pancreatic lipase activity were expressed as percentages of the control values which were obtained from the incubation of the reaction mixture containing only DMSO.

### RNA isolation and RT-PCR

Total RNA was extracted from tissues using the RNeasy kit (Qiagen, Hilden, Germany) and reverse transcribed using the TaKaRa PrimeScript RNA Kit (cat# RR036A, Takara, Japan), according to the manufacturer’s instructions. Quantitative PCR was performed using the TaqMan method (1 cycle at 50 °C for 2 min, 95 °C for 10 min, 40 cycles at 95 °C for 15 s, and 60 °C for 1 min) or the SYBR Green method (1 cycle at 95 °C for 30 s, 45 cycles at 95 °C for 10 s, and 60 °C for 20 s). Each sample was run in duplicate, and the relative mRNA levels were calculated using a standard curve and normalized to the mRNA levels of β-actin or GAPDH. The primer sequences used are listed in the Supplementary Table.

### 16S rRNA sequencing analysis

DNA was extracted from mouse cecal contents or feces using the QIAmp PowerFecal DNA kit (QIAGEN, CA, USA). A multiplexed amplicon library converting the 16S rDNA V4 region was generated from the DNA samples, and sequencing was performed in the Bioengineering Lab. Co., Ltd. (Kanagawa, Japan). Principal component analysis (PCA) was performed using the prcomp command in R version 3.2.1. The database consisted of Greengene’s 97 operational taxonomic units attached to the microbiota analysis pipeline, Qiime.

### Histological and Immunohistochemical Analysis

Sections of the liver, epididymal white adipose tissue (eWAT), and colon were excised and immediately fixed in 4 % formaldehyde at room temperature. Paraffin-embedded tissue sections were cut into 4 μm slices and placed on slides. Sections were stained with H&E or Alcian blue, according to standard procedures. Anti-CD36 antibody was purchased from Abcam (Cambridge, CB2 0AX, UK).

### Metabolome analysis (Dual Scan)

Metabolome analysis was conducted according to the HMT Dual Scan package using capillary electrophoresis time-of-flight MS and liquid chromatography time-of-flight MS in Human Metabolome Technologies Inc. (Japan).

### Western blotting

Proteins were extracted using 1× radioimmunoprecipitation assay buffer containing 0.1 % sodium dodecyl sulphate. Protein samples (18 μg) were subjected to SDS-PAGE and transferred to polyvinylidene fluoride membranes. Primary antibodies for β-actin (1:3000) were purchased from Cell Signaling Technology (Danvers, MA), claudin-1 was purchased from Invitrogen (1:200), and CD36 was from Abcam (1:100). Horseradish peroxidase-conjugated secondary antibodies were purchased from GE Healthcare (Japan) (1:1000). The intensities of the bands were detected using a ChemiDoc Touch MP system (Bio-Rad). ImageJ software was used for quantification.

### Statistical analyses

Statistical analyses were performed using GraphPad Prism 9 software (version 9.4.1; GraphPad Software, San Diego, CA, USA). Data are expressed as the means ± standard error of mean. *P < 0.05, **P < 0.01, ***P < 0.001, and ****P < 0.0001 were considered significant as determined using the unpaired two-tailed *t* test or analysis of variance (ANOVA) followed by Sidak’s multiple comparison test or Tukey-Kramer post hoc. Each dot represents a biological sample.

## Results

### IX suppresses body weight gain and improves glucose metabolism in mice on an HFD

HFD-fed C57BL/6 mice were fed a HFD supplemented with IX at 0.1%. IX significantly decreased body weight (Fig. 1A) without altering food intake (Fig. 1B). OGTT and ITT confirmed improved glucose metabolism (Fig. 1C and D), which was associated with decreased liver weight and improved steatosis (Fig. 1E and F). Increased cecum size was observed in the HFD+IX group, suggesting an altered intestinal environment after the intervention (Fig. 1E). Consistent with the altered liver weight, IX administration significantly reduced hepatic and plasma TG levels (Fig. 1G and H). Crown-like structures in the eWAT clearly decreased (Fig. 1I). The expressions of M1 macrophage related genes such as *Cd11c* and *Tnfα* were downregulated and metabolically favorable markers such as *Pgc1−a*, *Pgc1-b*, *Pparg*, and *adiponectin* were upregulated in the eWAT of IX group (Fig 1J). Next, we examined the effect of IX on thermogenesis-related genes such as *Ucp-1*, *Pgc1-a*, and *Cidea* in the inguinal adipose tissues which were unaffected by IX treatment (Suppl Fig. 1). Thus, IX administration improved glucose metabolism and decreased body weight gain in HFD-fed mice.

**Figure 1.**
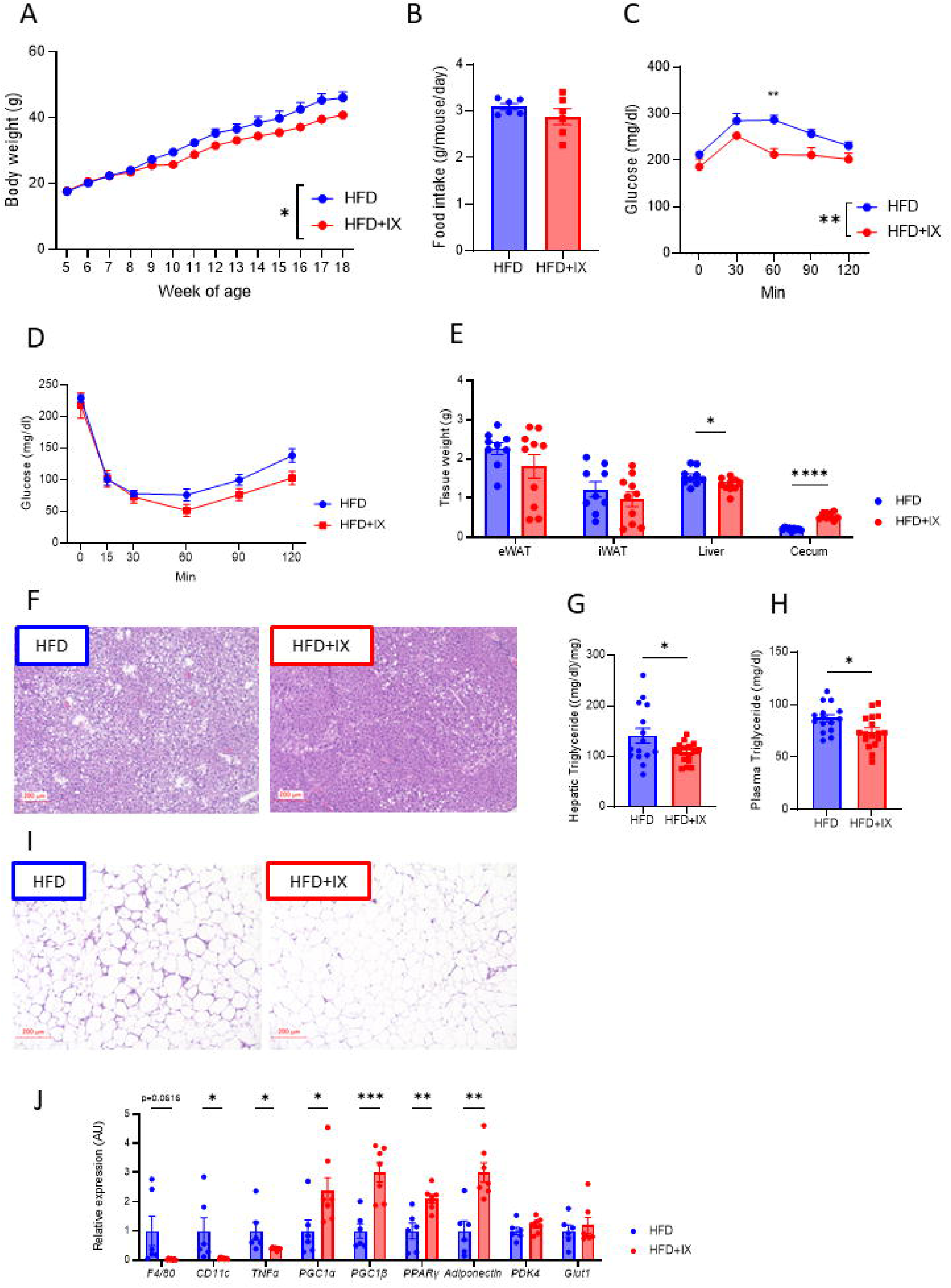
Isoxanthohumol (IX) suppresses body weight gain and improves glucose metabolism in mice on a high-fat diet (HFD). (A) Body weight and (B) daily food intake of mice treated with either an HFD (blue) or an HFD + IX (red) (n = 9–10 per group). (C) Oral glucose tolerance test (OGTT) at 17 weeks old and (D) insulin tolerance test (ITT) at 16 weeks old. (E) Tissue weight at 20 weeks old (n = 9–10 per group). (F, I) Representative hematoxylin and eosin (H&E)-stained pictures of (F) liver and (I) epididymal adipose tissue at 20 weeks old. Scale bars, 200 μm. (G) Hepatic and (H) plasma triglyceride concentrations at 20 weeks old (n = 15–17 per group). (J) Quantitative PCR analysis of inflammatory and metabolic markers in the epididymal adipose tissues at 20 weeks old (n = 6–7 per group). *P<0.05, **P<0.01, ***P<0.001, ****P<0.0001, by two-way analysis of variance (ANOVA) followed by Sidak’s multiple comparison test (A, C, D) or unpaired two-tailed *t* test (B, E, G, H, J). Data are presented as the mean ± SEM.

### IX pharmacologically inhibits lipid absorption

To further elucidate the mechanism of anti-obesity effect of IX administration, the amount of energy in the feces was analyzed with a bomb calorimeter. IX increased energy contents in the feces (Fig. 2A). Furthermore, we estimated the fecal lipid excretion by determining the lipid contents. The amounts of many types of fatty acids found in the NEFA and TG fractions from the fecal lipids were significantly increased by IX, suggesting that IX inhibited lipid absorption (Fig 2B and C).

**Figure 2.**
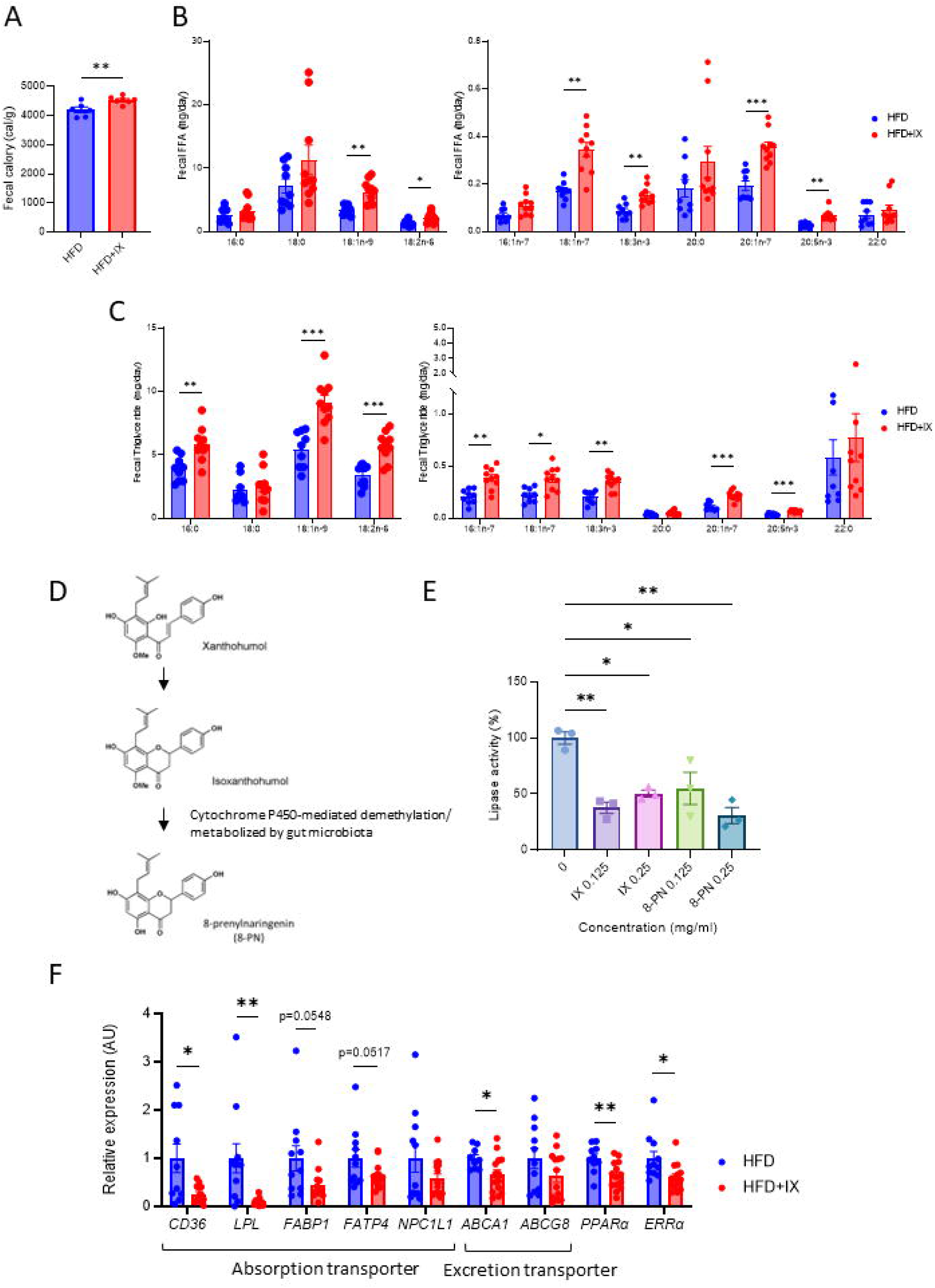
IX promotes intestinal lipid excretion. (A) Amount of energy in feces collected for 24 h from each mouse (n = 6-7 per group). (B, C) Concentrations of various classes of (B) fatty acids and (C) triglycerides in feces collected at 24 h from each mouse (n = 9–10 per group). (D) A simplified metabolic pathway of xanthohumol in the gut microbiota. (E) Lipase activity of IX and 8-prenylnaringenin at the indicated concentrations (n = 3 per group). (F) qPCR analysis of various lipid transporter-related genes in the jejunum of mice at 18 weeks old (n = 11-14 per group). *P<0.05, **P<0.01, ***P<0.001, by unpaired two-tailed *t* test for (A, B, C and F) and ANOVA, followed by Tukey-Kramer post doc for (E). Data are presented as the mean ± SEM.

IX is known to be converted to 8-prenylnaringenin (8-PN) via hepatic cytochrome P450-mediated demethylation or can be metabolized by gut bacteria (Fig 2D) [17]. To explore the mechanism underlying the inhibition of fatty acid absorption by IX, we assessed the lipid digestive capacity and found that, both, IX and 8-PN inactivated pancreatic lipase activity by approximately 50–60 % (Fig. 2E). On the other hand, the expression of *Cd36*, a major lipid transporter was significantly downregulated in line with decreased lipoprotein lipase, *Ppara,* and *Erra* expressions in the small intestine of mice treated with HFD+IX (Fig. 2F). These data indicate that IX or its metabolite 8-PN inhibit the breakdown of dietary TGs and lower absorption of fatty acids from the small intestine.

Further, we tested whether IX or 8-PN directly regulates fatty acid transporter gene expression in cultured intestinal epithelial cells (Supplementary Fig. 2A). The expression of *Cd36*, *Fabp1*, and *Fatp4* was not altered by the treatment of isolated epithelial cells with IX or 8-PN for 24 h (Supplementary Fig. 2B). Thus, IX and 8-PN do not directly regulate the expression of fatty acid transporter-related genes in the intestinal epithelial cells.

We also performed a plasma metabolomic analysis to identify the metabolites altered by IX administration. Heatmap and PCA did not reveal significant differences (Supplementary Fig. 3A, B). Volcano plots identified that IX treatment significantly altered a small set of seven metabolites (Supplementary Fig. 3C) among two of which were phytosterols, sitosterol and campesterol (Supplementary Fig. 3C). To determine the metabolic effects of these two phytosterols that were decreased by IX treatment, the mice were treated with phytosterol for 11 weeks. This had no effect on the body weight of mice fed either chow diet or HFD (Supplementary Fig. 3E). On chow diet, phytosterol administration increased fasting blood glucose, but did not affect glucose tolerance on chow or HFD (Supplementary Fig. 3F). Thus, the reduction of plasma phytosterols by IX administration had no significant direct effect on obesity or glucose metabolism.

### The favorable metabolic effects of IX treatment are nullified by the elimination of gut microbiota

Cecal weight significantly increased in in IX-treated mice (Fig. 1E). Since the cecum size can reflect altered gut microbiota [18], we evaluated the metabolic effects of IX treatment without the influence of the microbiota. Interestingly, eliminating gut microbiota by antibiotic treatment nullified the IX-induced metabolic improvements, such as body weight, glucose metabolism, and tissue weight (Fig. 3A-D). Antibiotic treatment resulted in an even greater cecal weight, reflecting a marked decrease in microbial content in the intestine (Fig. 3D), in agreement to what is known for GF mice[18]. Fecal TG content was elevated in the IX group, which significantly decreased after antibiotic treatment (Fig. 3E). These results highlight the favorable metabolic effects of IX treatment only in the presence of intact gut microbiota.

**Figure 3.**
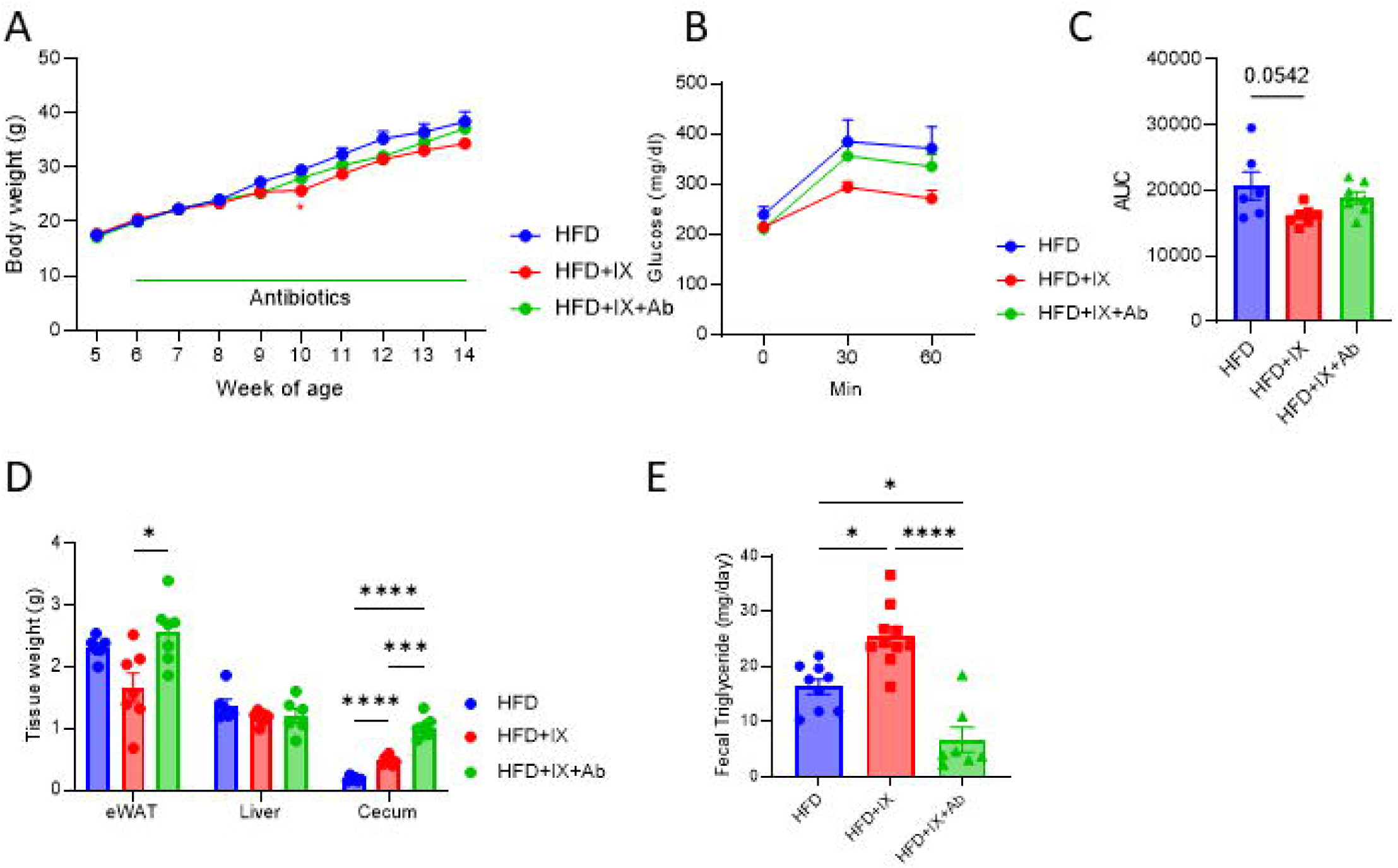
The gut microbiota established by IX improves metabolic dysfunction. (A) Body weight of mice treated with either an HFD (blue), an HFD + IX (red) or an HFD + IX + antibiotics (green) (n = 6–7 per group). (B) OGTT at 15 weeks old and (C) area under the curve (AUC) measured during OGTT (n = 6–7 per group). (D) Tissue weight at 18 weeks old (n = 6–7 per group). (E) Fecal triglyceride concentrations (n = 7–10 per group). *P<0.05, **P<0.01, ***P<0.001, ****P<0.0001, using ANOVA, followed by Tukey-Kramer post doc for (C, D, E). Data are presented as the mean ± SEM.

To determine the effects of the microbiota established by IX treatment, 8-week-old GF mice were colonized with the microbiota from either HFD-or HFD+IX-fed mice for 5 weeks (Supplementary Fig. 4A). The cecum of the recipient mice given HFD+IX was enlarged, reflecting the donor phenotype, suggesting the viability of the transplanted bacteria (Supplementary. Fig. 4B). Although the effect of FMT from HFD+IX donors on liver weight was not altered at 13 weeks age (Supplementary. Fig 4B), body weight and glucose tolerance tended to improve transiently (Supplementary. Fig. 4C and D). These metabolic changes were relatively minor, probably because the animals were fed a chow diet.

**Figure 4.**
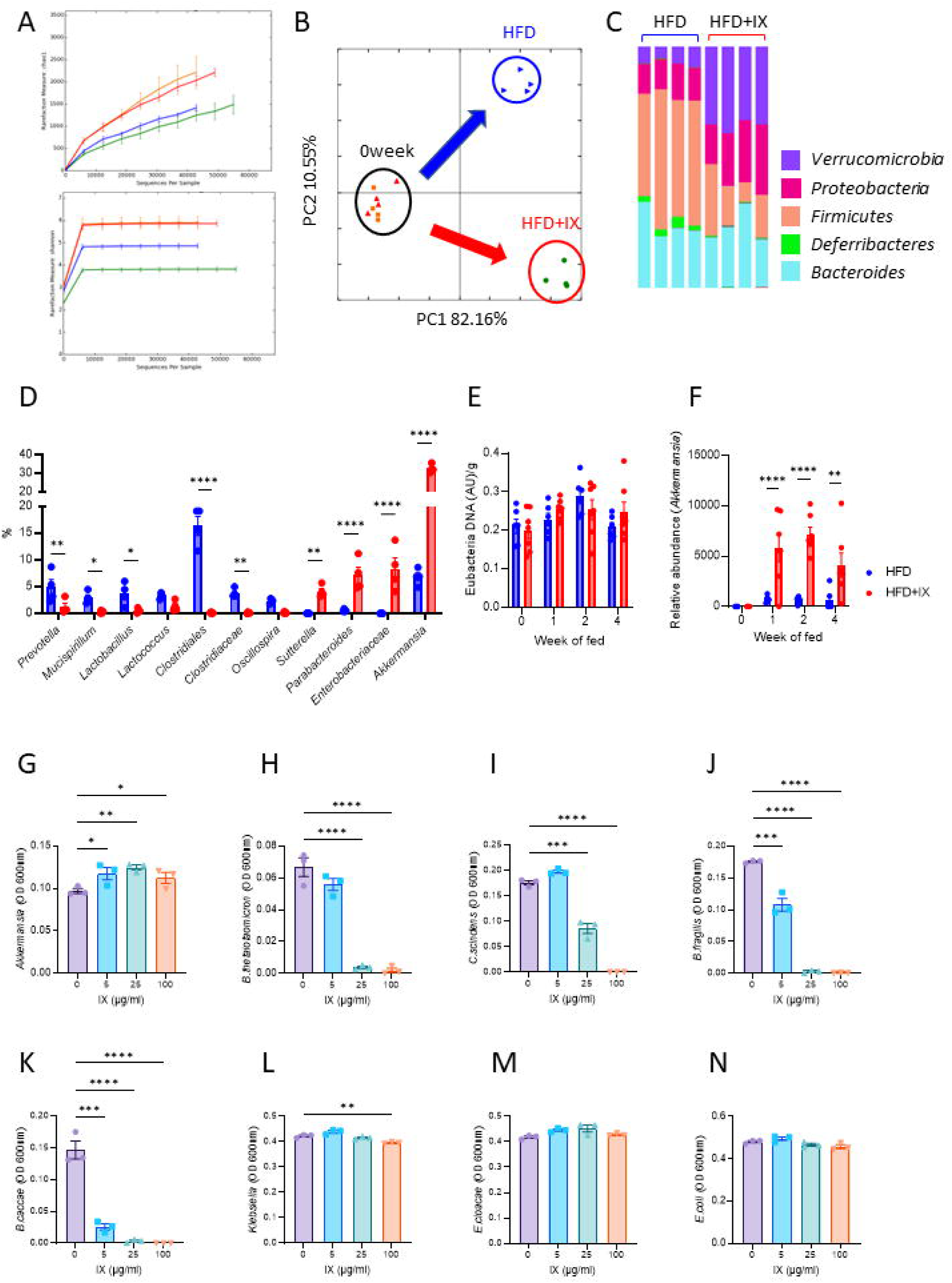
IX promotes the growth of *A. muciniphila*. (A)Rarefaction curves of Chao1 and Shannon entropy of fecal 16S rRNA sequencing data from HFD-fed mice with or without IX for 2 weeks (red: pre-HFD, blue: post-HFD orange: pre-HFD+IX, green: post-HFD+IX). (B) Principal coordinate analysis of weighted UniFrac distances. (C) Representation of bacterial phyla in the fecal bacteria of HFD-fed mice with or without IX for 2 weeks. (D) Relative abundances of bacteria that showed significant differences between the HFD and HFD + IX groups (n = 4 per group). (E) Eubacterial DNA levels per gram of feces at different time points (n = 6–7 per group). (F) Relative abundance of *A. muciniphila* normalized to eubacterial levels (n = 6–7 per group). (G)–(N) Growth of bacteria as single cultures in the presence or absence of IX at the indicated concentrations. *P<0.05, **P<0.01, ***P<0.001, ****P<0.0001, by unpaired two-tailed *t* test for (D, E and F) and ANOVA, followed by Tukey-Kramer post dot for (G-N). Data are presented as the mean ± SEM.

### IX impacts the microbial community structure and promotes the growth of *A. muciniphila*

Rarefaction curves obtained from 16S rRNA sequencing analysis of fecal samples revealed IX-induced reduction in species richness and evenness (Fig. 4A), suggesting the predominance of certain bacterial species. Principal coordinate analysis of weighted UniFrac distances showed a clear separation in the microbial community structures before and after the HFD or HFD + IX challenge (Fig. 4B). IX increased the relative abundance of the phylum Verrucomicrobia and decreased those of Firmicutes and Deferribacteres (Fig. 4C). At the lower phylogenetic tree level, various bacteria, such as *Mucispirillum*, *Lactobacillus*, Clostridiales, and Clostridiaceae, were eliminated after IX treatment (Fig. 4D). In contrast, some bacteria classified as *Parabacteroides*, *Sutterella*, *Enterobacteriaceae*, and *A. muciniphila* were elevated in the IX group (Fig. 4D). Since *A. muciniphila* has beneficial effects on glucose metabolism and was most significantly increased by IX treatment, we followed the time course of the quantitative changes after the intervention. While eubacterial DNA levels, which reflect the total bacterial biomass, were not altered by IX treatment for at least the first 4 weeks, the relative abundance of *A. muciniphila* markedly increased after 1 week (Fig. 4E and F).

To investigate mechanisms for the proliferation of *A. muciniphila*, we evaluated the effects of IX on bacterial proliferation using an anaerobic chamber. The *in vitro* study showed that IX selectively promoted the growth of *A. muciniphila* in pure cultures, but not that of other bacteria examined (Fig. 4G-N). Thus, IX significantly altered the microbial community structure, directly promoting the growth of *A. muciniphila* and inhibiting the growth of other bacteria, thereby causing a proliferation of *A. muciniphila* in the intestine.

### IX strengthened the gut barrier function

Interventions to increase the abundance of *A. muciniphila* improve intestinal barrier function[10]. Hence, we further examined whether the intestinal environment, with IX-induced proliferation of *A. muciniphila*, enhanced barrier function. IX administration thickened the mucus layer of the colon (Fig. 5A), which was confirmed by the quantification of fecal mucin levels (Fig. 5B). Tight junction-related claudin 1 level was also increased in the IX group (Fig. 5C-D), which was associated with the decreased expression of lipopolysaccharide binding protein (LBP), an endotoxin marker, in the liver (Fig. 5E). Thus, IX improved the colonic barrier function through the proliferation of *A. muciniphila* to improve insulin sensitivity.

**Figure 5.**
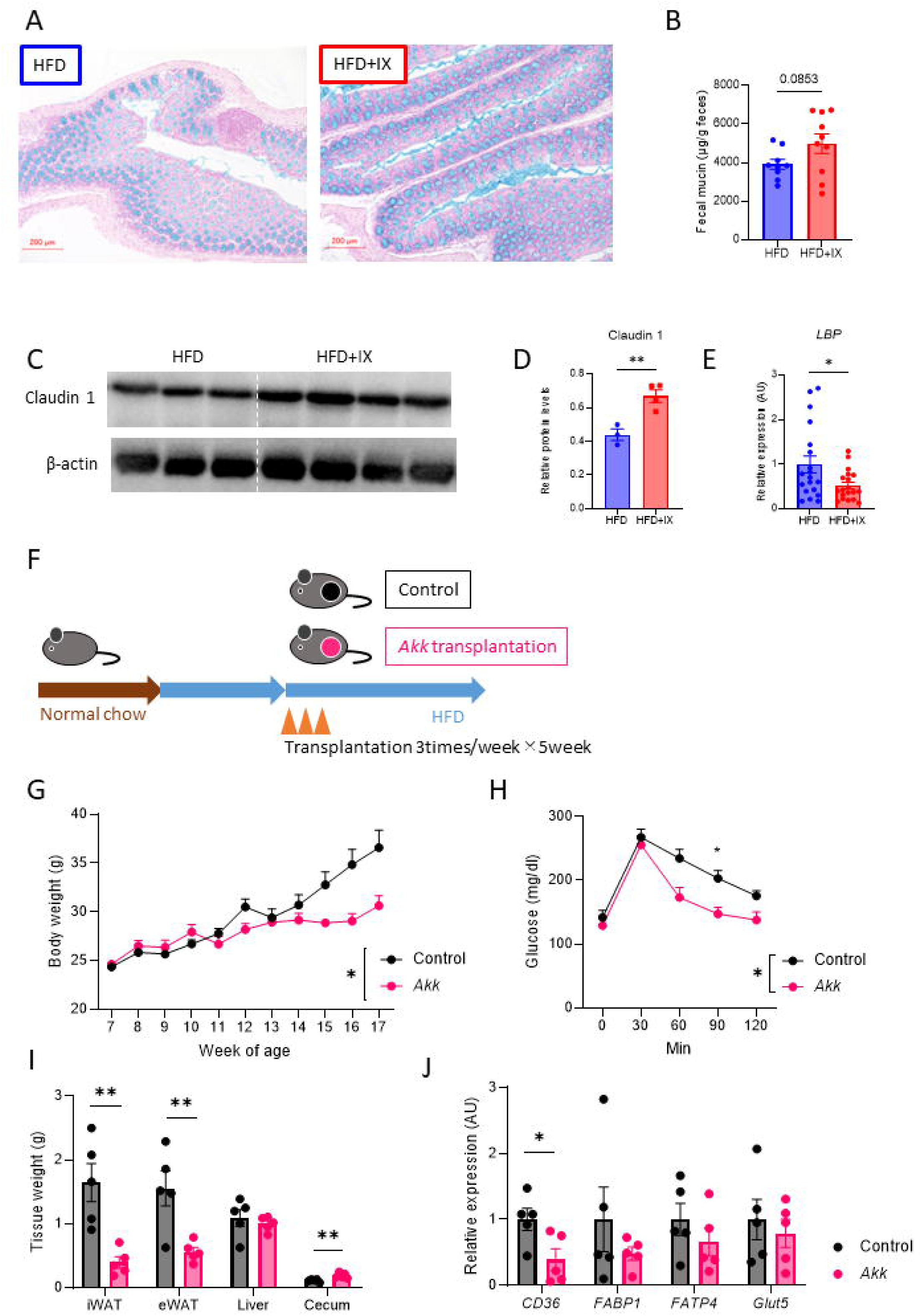
Compound IX enhances gut barrier function. (A) Representative alcian blue images of the colon. Scale bars, 200 μm. (B) Fecal mucin levels per gram feces of mice fed either an HFD or an HFD+IX (n = 9–10 per group). (C) Western blots for claudin1 and β-actin in the colon of mice at 20 weeks old. (D) Quantitation of claudin1 normalized by β-actin. (E) qPCR of the lipopolysaccharide binding protein in the liver (n = 19 per group). (F) Schematic overview of the transplantation of specific pathogen-free mice with *A. muciniphila*. (G) Body weights of transplanted mice (n = 5 per group). (H) OGTT. (I) Tissue weight. (J) qPCR of various nutrient transporter-related genes in the jejunum of the transplanted mice (n = 5 per group). *P<0.05, **P<0.01, using two-way ANOVA, followed by Sidak’s multiple comparison test for (G, H) or by unpaired two-tailed *t* test for (B, D, E, I, J). Data are presented as the mean ± SEM.

### *A. muciniphila* colonization inhibits fatty acid absorption to prevent obesity

We further examined the effects of orally administering *A. muciniphila* on fat accumulation in HFD-fed mice. The mice were gavaged with *A. muciniphila* thrice a week for 5 weeks (Fig. 5F). The continuous administration of *A. muciniphila* markedly suppressed weight gain and improved glucose tolerance (Fig. 5G and H). These effects were associated with decreased adipose tissue size and *Cd36* downregulation in the jejunum (Fig. 5I and J), indicating that *A. muciniphila* inhibited fatty acid absorption and prevented fat accumulation.

To determine the direct effects of *A. muciniphila* on energy and glucose metabolism, we colonized GF mice with either *A. muciniphila* or *B. thetaiotaomicron,* which reside in the mucus layer (Fig. 6A). After 1 week of monocolonization on a normal diet, increased body and liver weights were observed among mice in the *B. thetaiotaomicron* group compared to those in GF mice, while *A.muciniphila* monocolonization had no effect on body or liver weights (Fig. 6B and C). The OGTT revealed improved glucose tolerance in the *A. muciniphila* group compared to that in the *B. thetaiotaomicron* group (Fig. 6D-E). *A. muciniphila* specifically suppressed the expression of *Cd36*, the main fatty acid transporter among the various nutrient transporters (Fig. 6F and G). Finally, we performed plasma metabolomic analysis and found that *A. muciniphila* colonization lowered the plasma levels of various classes of fatty acids (Fig. 6H). In summary, IX induced *A. muciniphila* proliferation, which enhanced intestinal barrier function and inhibited fatty acid absorption from the small intestine, leading to reduced obesity and improved insulin resistance.

**Figure 6.**
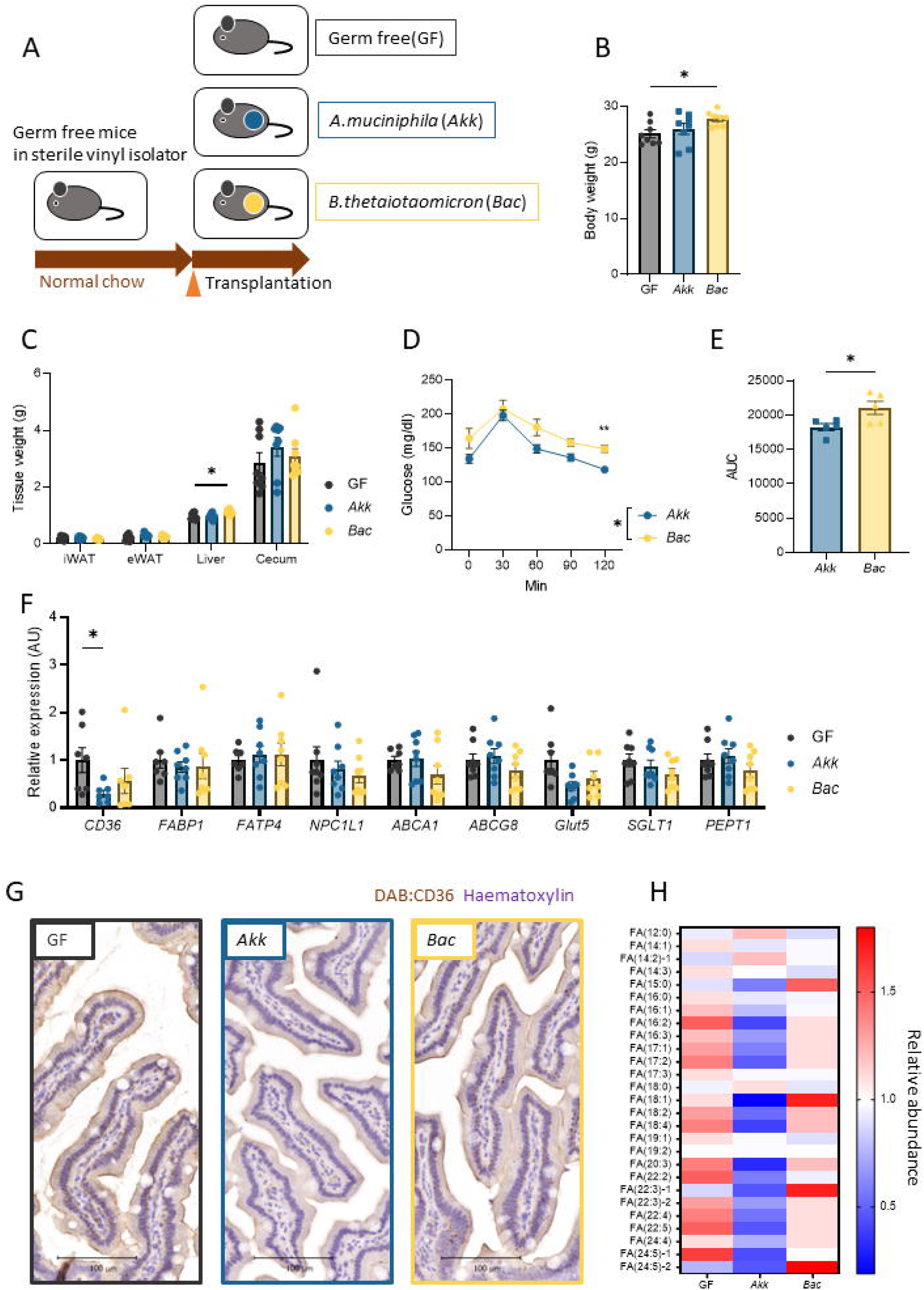
*A. muciniphila* suppresses fatty acid absorption via decreasing transporter-related genes in the small intestine. (A) Schematic overview of monocolonization of germ-free mice with *A. muciniphila* or *Bacteroides thetaiotaomicron*. (B) Body weight of monocolonized mice (n = 8 per group). (C) Tissue weight (D) OGTT (E) AUC of OGTT (F) qPCR of various nutrient transporter related genes in the jejunum of monocolonized mice (n = 8 per group). *P<0.05, **P<0.01, using two-way ANOVA, followed by Sidak’s multiple comparison test for (C, D), or using ANOVA, followed by Tukey-Kramer post doc for (B, E, F). Data are presented as the mean ± SEM. (G) Representative CD36 images of the jejunum. Scale bars, 100 μm. (H) Heatmap showing saturated or unsaturated fatty acids detected in the plasma of monocolonized mice.

**Figure 7.**
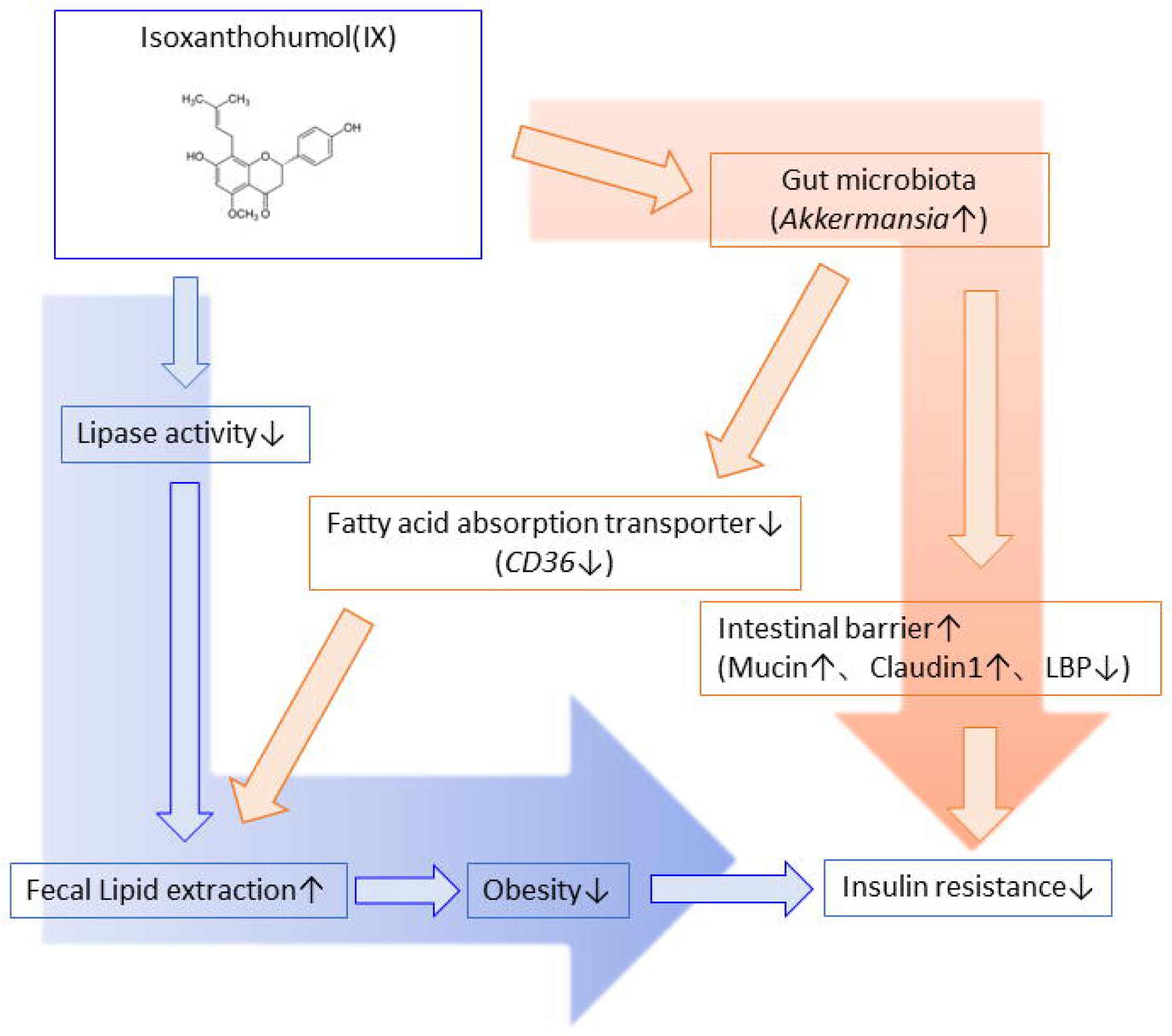
Proposed mechanisms of IX on obesity and insulin resistance. As a pharmacological pathway, IX inhibits lipase activity and reduces the expression of *Cd36* in the small intestine, thereby increasing fecal lipid excretion and obesity. In the microbial pathway, IX improves insulin resistance by altering the microbial composition, specifically increasing the abundance of *A. muciniphila* and enhancing intestinal barrier function. The anti-obesity effect of IX is further enhanced by the inhibition of fat absorption promoted by the proliferation of *A. muciniphila*.

## Discussion

Disruption of the gut microbiota significantly impacts energy metabolism. Turnbaugh et al. discovered that the gut microbiota in obesity more efficiently harvest dietary energy, further inducing obesity[2, 3]. Thus, manipulating gut microbiota can potentially contribute to treating obesity. Various actions of microbial metabolites regulating energy metabolism have been elucidated. SCFAs and the activation of TGR5 regulate the energy balance towards anti-obesity[4]. However, strategies to modulate the nutrient absorption capacity of the intestine, including the microbiota, are still unclear.

Polyphenols in hops, a beer ingredient, are beneficial to health[12]. Xanthohumol inactivates sterol regulatory element-binding proteins and reduces fatty acid synthesis, thus regulating lipid metabolism and improving obesity[19]. It also possesses biological activities which improve glucose metabolism in the presence of the gut microbiota[20]. This corroborates the supposition that the ingredients in beer possess antimicrobial properties [21].

IX is a heat-stable prenylflavonoid produced from xanthohumol during beer brewing [22], which has been suggested to improve obesity and insulin resistance [15, 23], but the mechanism has not been clarified yet. A possible mechanism involves microbial metabolites, such as SCFAs, but IX administration does not alter fecal SCFA levels [24]. In this study, we performed plasma metabolomic analysis of IX-treated mice and found no significant difference in SCFA levels in the IX treated-mice compared to those in the control mice, suggesting that the metabolic improvement effect promoted by IX was independent of SCFAs.

Here, IX treatment improved obesity and metabolic disorders through the inhibition of pancreatic lipase activity and intestinal bacteria-mediated action. The inhibition of pancreatic lipase reduces lipid absorption by attenuating the hydrolysis of dietary TGs. Orlistat, a lipase inhibitor, similarly inhibits lipid absorption and exerts its weight-loss effect; however, the adverse effects related to gastrointestinal symptoms such as diarrhea and steatorrhea limit its extensive use. [25]. While administering 0.012 % orlistat to an HFD increases the stool TG content by approximately 176 % from 151 mg/g feces to 417 mg/g feces[26], IX administration increased it by 44 % (Fig. 3E). Thus, the lipid excretion-promoting effect of IX and the related gastrointestinal symptoms appeared to be milder than those of orlistat.

IX treatment significantly altered the microbial community structure and the favorable metabolic effects were canceled by antibiotic treatment, suggesting an involvement of the gut microbiota. Some polyphenols increase a relative abundance of *A. muciniphila* and are beneficial for metabolic health[13, 27]. In the present study, IX selectively promoted the growth of *A. muciniphila* in an anaerobic chamber study, and significantly increasing the abundance of *A. muciniphila* in vivo.

*A. muciniphila* is a gram-negative anaerobic bacterium associated with ameliorating metabolic disorders in obese humans and rodents[11, 28]. Provier et al. showed that administering live or pasteurized *A. muciniphila* reduces weight gain, insulin resistance, and dyslipidemia; they proposed the importance of TLR2 activation via the membrane protein components from *A. muciniphila*[28]. Furthermore, pasteurized *A. muciniphila* exhibits anti-obesity properties by increasing systemic energy expenditure, as assessed by metabolic cages and fecal energy excretion. Their study revealed downregulation of carbohydrate transporter-related genes, which is associated with enhanced epithelial turnover in the jejunum[29]. These results differed from the inhibition of lipid absorption by IX and *A. muciniphila* in our study, possibly because they administered perturbed *A. muciniphila* and we intended to explore the direct action of a single bacterial species via monocolonizing *A. muciniphila.* However, this is consistent with the fact that *A. muciniphila* exhibits anti-obesity effects by regulating intestinal nutrient absorption.

*A. muciniphila* also enhances barrier function [10]. Despite being a mucin-degrading bacterium, *A. muciniphila* administration increases mucus thickness[30] and goblet cell density[28]. This effect may be promoted by the enhanced turnover of epithelial cells [29] or the production of SCFAs, which are a major nutrient source for intestinal cells[31]. Deterioration of the intestinal environment due to an HFD and obesity reduces the intestinal barrier function and causes insulin resistance through metabolic endotoxemia [32, 33]. Thus, IX-induced *A. muciniphila* proliferation restored the impairment of barrier function and improved insulin resistance, as confirmed by increased mucin and claudin-1 levels, and decreased LBP expression in the liver.

Furthermore, the *A. muciniphila* monocolonized mouse model and its metabolomic analysis revealed that *A. muciniphila* exhibits an anti-obesity effect by downregulating Cd36 expression in the small intestine and inhibiting fatty acid absorption. This is a novel mechanism underlying the effect of a single *A. muciniphila* strain on energy metabolism.

CD36 is a fatty acid transporter expressed in many cells and tissues, including platelets, macrophages, intestinal epithelial cells, endothelial cells, smooth muscle cells, adipose tissues, skeletal muscles, and cardiomyocytes. In the intestine, CD36 is highly expressed on the brush border membrane of enterocytes and is mainly localized in the duodenum and jejunum[34]. In addition, HFD significantly increases CD36 expression in the jejunum of mice[29, 35], and its expression positively correlates with increasing BMI. In addition, it is significantly increased in vascular lesions and the kidneys of patients with hyperglycemia and/or hyperlipidemia, suggesting the dysregulation of CD36 levels in obesity and related metabolic dysfunction[36]. Although CD36 has intestinal hormone modulatory effects on GIP, GLP-1, and secretin, it is a major regulator of lipid absorption, as evidenced by a 50 % reduction in lipid absorption in an intestinal *Cd36*-deficient mouse model[37]. C24:0 fatty acid absorption is completely inhibited from the intestine of *Cd36*-deficient mice fed an HFD, highlighting the important role of intestinal CD36 in absorbing dietary long-chain fatty acids[38].

Various signaling pathways regulate CD36 expression in enterocytes. Intestinal hormones such as secretin and cholecystokinin can act on their receptors to upregulate CD36 expression, promoting intestinal lipid absorption [39, 40]. In addition, diurnal variations in histone deacetylase 3, which is regulated by the microbiota, positively regulates CD36 via the activation of ERRα[41]. Stojanović et al. reported that CD36 is downregulated in *Ppara*-deficient mice, suggesting that PPARα is a key regulator of intestinal CD36[42]. In our study, IX treatment significantly decreased the expression of both *Ppara* and *Erra*, suggesting that IX regulates CD36 through these pathways.

As an interaction between intestinal bacteria and CD36 expression, FMT from HFD-fed mice into GF mice induces CD36 expression in the small intestine[43]. This suggests a close relationship between the adaptation of microbiota to nutrients and nutrient absorption. Kawano et al. reported that Th17-inducing microbiota, segmented filamentous bacteria (SFB) downregulates *Cd36* expression in the small intestine and prevents weight gain; however, *Faecalibaculum rodentium*, which is increased by a high-sugar diet, eliminates SFB and accelerates obesity[44]. Thus, intestinal bacterial interactions may be involved in CD36 regulation. To our knowledge, this is the first study to show that monocolonization of *A. muciniphila*, which was markedly increased by IX, decreases CD36 expression in the small intestine and plasma fatty acid concentrations using metabolomics analysis. Since HFD and obesity decrease the relative abundance of *A. muciniphila,* this may disrupt lipid absorption regulation, further contributing to obesity.

In conclusion, our study showed that IX prevents obesity and improves glucose metabolism by suppressing dietary fat absorption through suppressing pancreatic lipase activity and the microbial action by selectively promoting the growth of *A. muciniphila* and improving intestinal barrier function. Furthermore, its anti-obesity effect is partly explained by the inhibition of fatty acid absorption due to *A. muciniphila*-induced reduction of the fatty acid transporter CD36 in the small intestine. Response of the microbiota to food components can control intestinal function and energy metabolism, suggesting a potential new therapeutic strategy against obesity.

## Supporting information

Primer

## Data Availability

The datasets generated and analyzed in this study are available from the corresponding author on reasonable request.

## Author contributions

Y.W. and S.F. designed and performed the experiments, analyzed the data, and wrote the manuscript. Y.M. helped bacterial experiments. S.W. performed the biochemical analyses. K.H. contributed technical assistance. H.H. performed histological experiments. A.N., To.K., Ay.N, M.B., A.R., Y.N., S.S., K.H., T.N., Y.N. and K.T. contributed to the discussion and interpretation of the data.

## Acknowledgments

We thank Yurie Iwakuro and Kumi Sassa for their technical assistance.

## Abbreviations

8-PN: 8-prenylnaringenin
BHI: Brain Heart Infusion
ELSD: evaporative light-scattering detection
eWAT: epididymal white adipose tissue
FMT: Fecal microbiota transplantation
GF: Germ-free
GLP-1: glucagon-like peptide 1
HFD: high-fat diet
IX: Isoxanthohumol
NEFA: non-esterified fatty acid
PCA: Principal component analysis
PCA: Principal component analysis
SCFA: short-chain fatty acid
TG: triglyceride
TGR5: G-protein-coupled bile acid receptor

---

**Supplementary Figure1.**
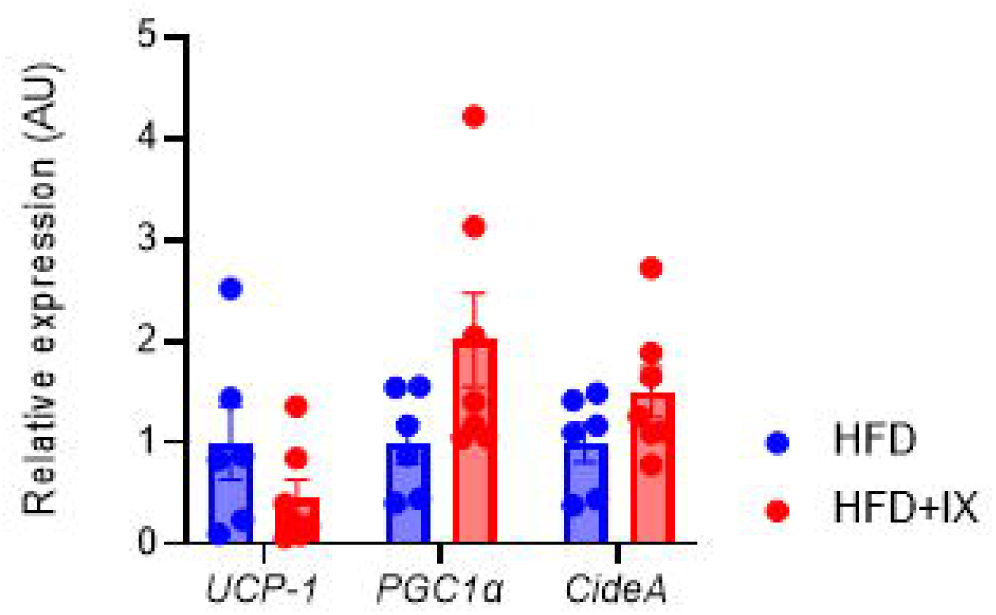
JX administration hadno influence on thermogenesis in the inguinal adipose tissue. qPCR analysis of thermogenesis related markersin the inguinal adipose tissues at 20 weeks old (n =6-7 per group).

**Supplementary Figure 2.**
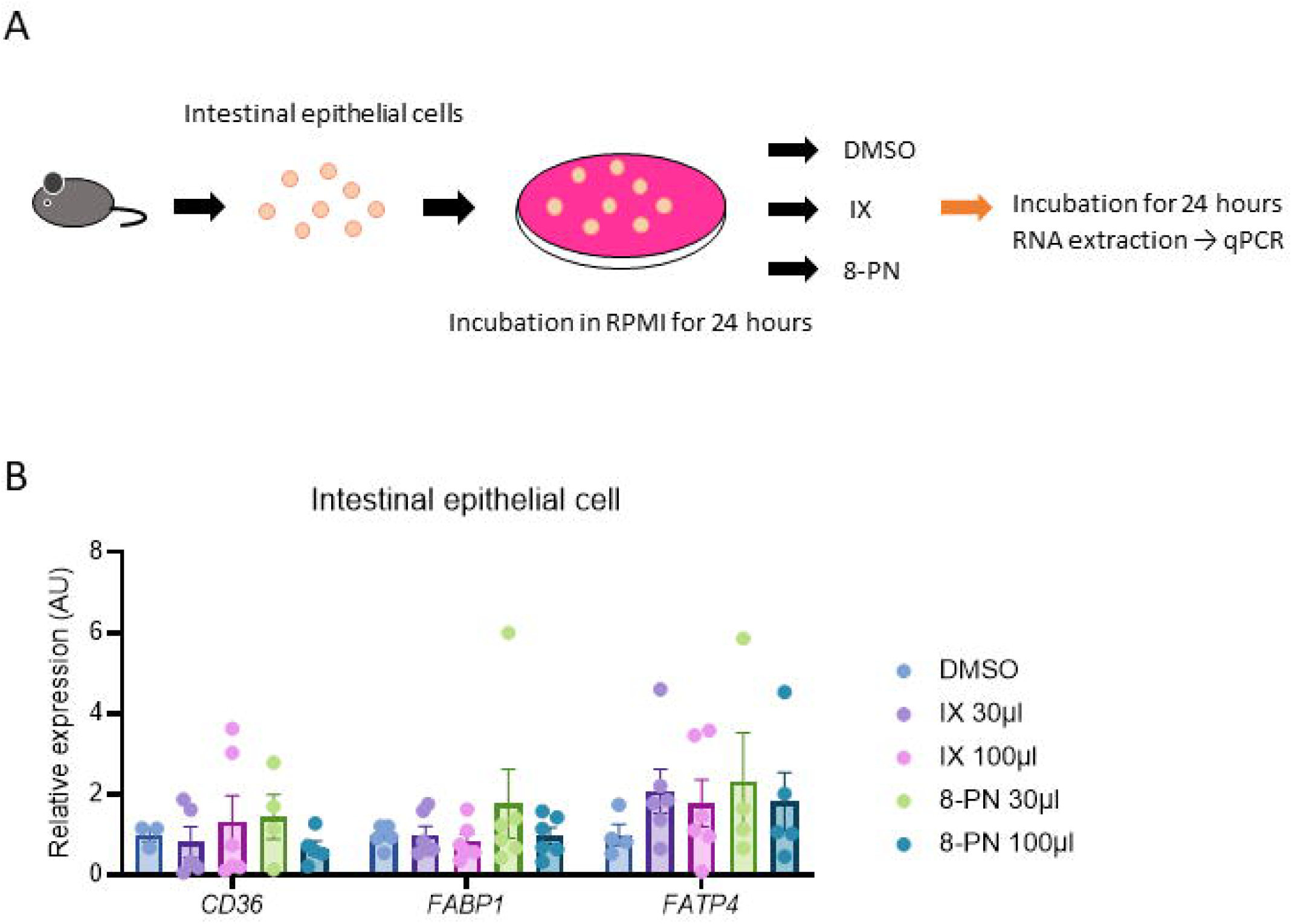
IX and 8-PN haveno direct function on fattyacid transportersin the intestinal cells. (A) Schematicoverview of incubation of the intestinal epithelial cells.(B) qPCRanalysisof fatty acid absorption related markersin the intestinal epithelial cells.

**Supplementary Figure 3.**
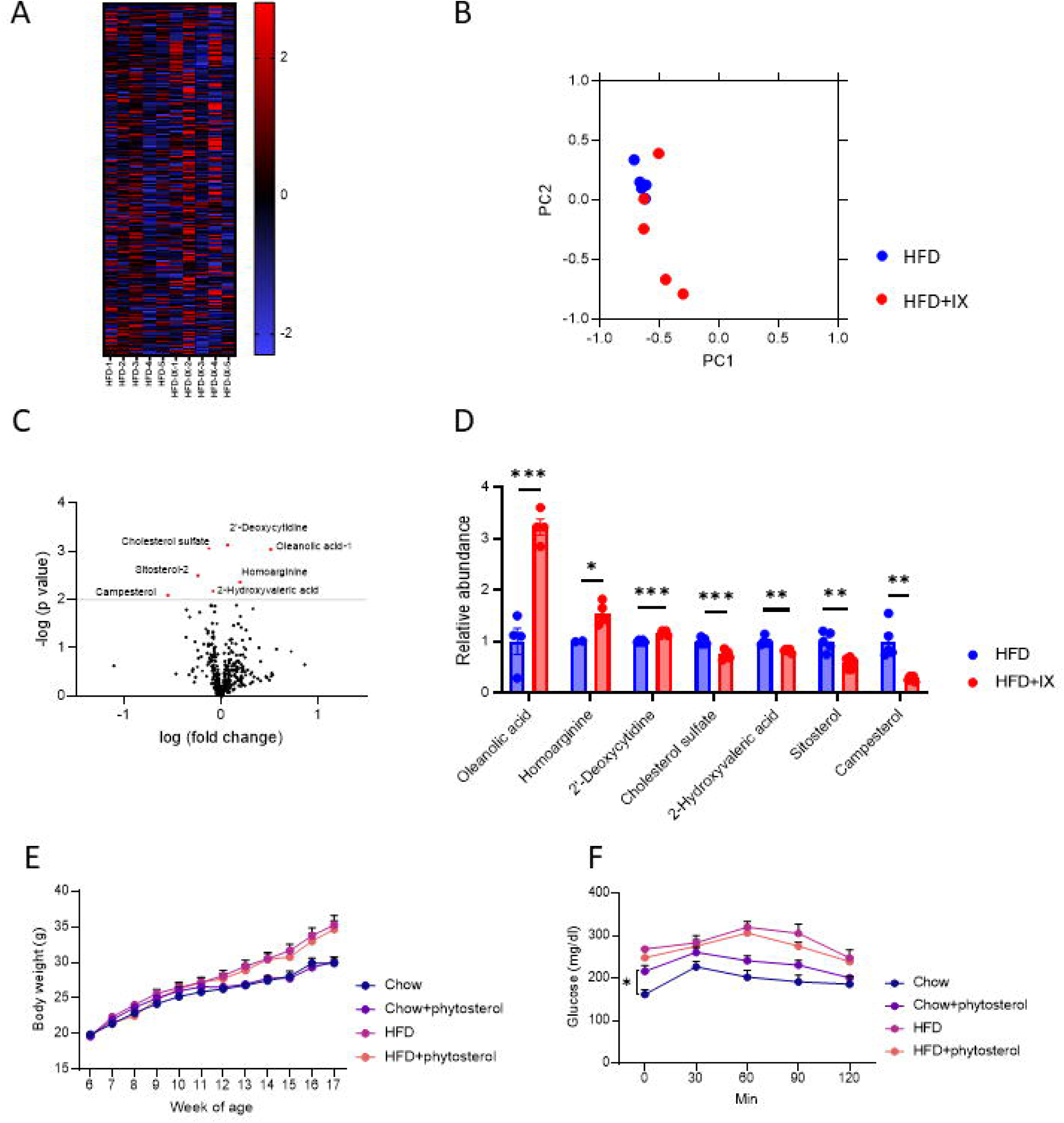
Metabolites derived fromIX administrationhave small impact on obesity or glucose metabolism of mice. {A-0)Heatmap {A), PCA analysis{B), volcano plot (C) of metabolomics of mice at18 weeks old (n =5 per group).(D) Metabolitesshows significant change withIX administration.{E-F) Body weight(E) and oral glucose tolerance testat17 weeksage {F) of mice fed chow,chow withphytosterol,HFD,orHFD withphytosterol (n = 5-6 per group).Phytosteroladministration was startedat 6 weeks age. *P<0.05, **P<0.01, ***P<0.001, by unpaired ttest{D), or by two-way ANOVA followed byTukey-Kramerpostdoc(E, F).

**Supplementary Figure 4.**
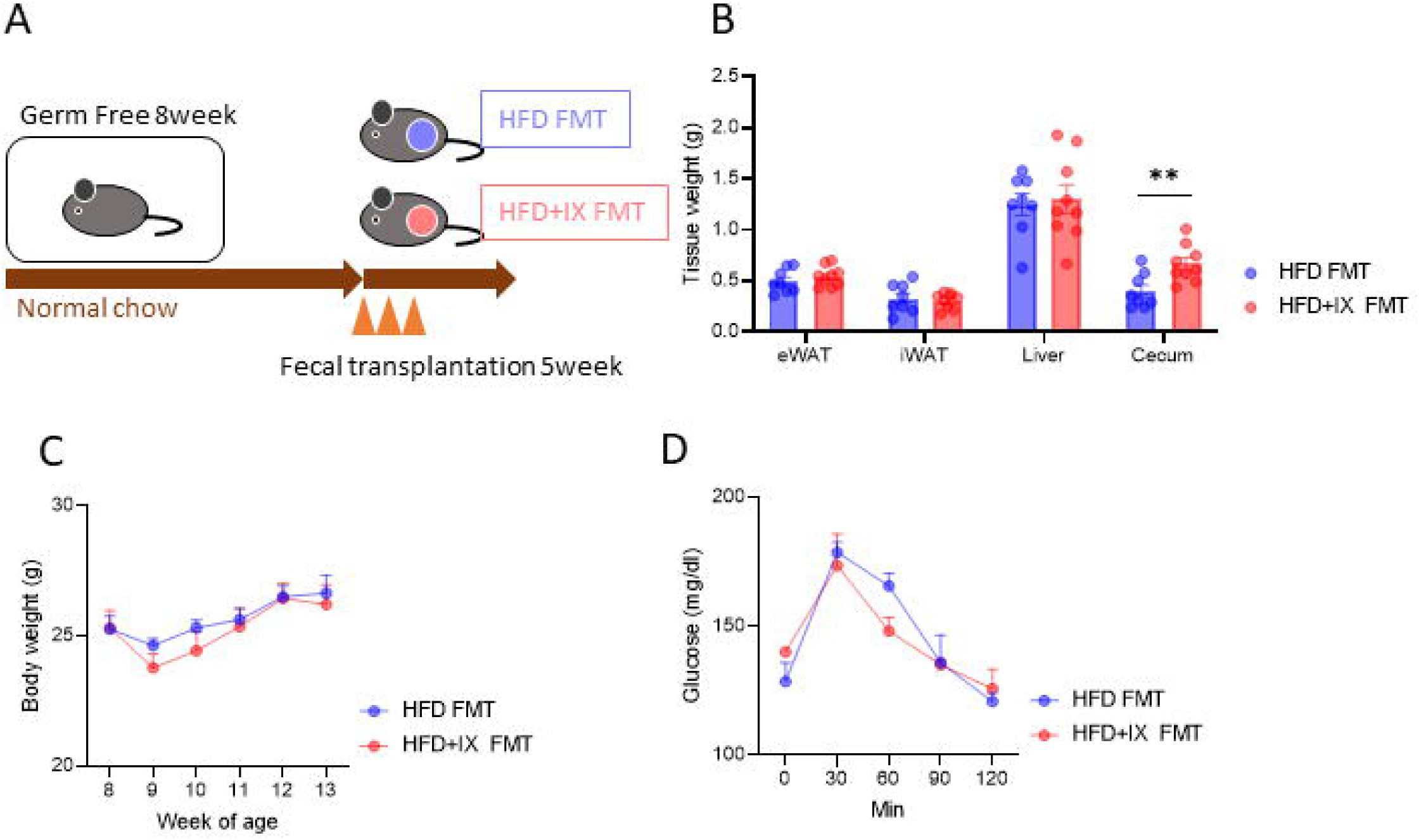
Effectsof FMT from HFO+IX mice to GF mice wererelatively minerunderchow-fed condtion. (A) Schematicoverview of the gnotobiota study. (B)Tissue weight(C)Body weight of the recipient mice transferred fecal microbiota fromeither anHFO or an HFO+IX fed mice(n = 8-9 per group).(0) OGTT performed 5 weeksafter transplantation(n =8-9 per group). **P<0.01 by unpaired*ttest.*Data are presented as the mean ± SEM.

## References

[1] Koh A, Backhed F (2020) From Association to Causality: the Role of the Gut Microbiota and Its Functional Products on Host Metabolism. Mol Cell 78(4): 584–596. 10.1016/j.molcel.2020.03.005

[2] Ridaura VK, Faith JJ, Rey FE, et al. (2013) Gut microbiota from twins discordant for obesity modulate metabolism in mice. Science 341(6150): 1241214. 10.1126/science.1241214

[3] Turnbaugh PJ, Ley RE, Mahowald MA, Magrini V, Mardis ER, Gordon JI (2006) An obesity-associated gut microbiome with increased capacity for energy harvest. Nature 444(7122): 1027–1031. 10.1038/nature05414

[4] Fujisaka S, Watanabe Y, Tobe K (2023) The gut microbiome: a core regulator of metabolism. J Endocrinol 256(3). 10.1530/JOE-22-0111

[5] Kimura I, Ichimura A, Ohue-Kitano R, Igarashi M (2020) Free Fatty Acid Receptors in Health and Disease. Physiol Rev 100(1): 171–210. 10.1152/physrev.00041.2018

[6] van Nierop FS, Scheltema MJ, Eggink HM, et al. (2017) Clinical relevance of the bile acid receptor TGR5 in metabolism. Lancet Diabetes Endocrinol 5(3): 224–233. 10.1016/S2213-8587(16)30155-3

[7] de Vos WM, Tilg H, Van Hul M, Cani PD (2022) Gut microbiome and health: mechanistic insights. Gut 71(5): 1020–1032. 10.1136/gutjnl-2021-326789

[8] Derrien M, Vaughan EE, Plugge CM, de Vos WM (2004) Akkermansia muciniphila gen. nov., sp. nov., a human intestinal mucin-degrading bacterium. Int J Syst Evol Microbiol 54(Pt 5): 1469–1476. 10.1099/ijs.0.02873-0

[9] Dao MC, Everard A, Aron-Wisnewsky J, et al. (2016) Akkermansia muciniphila and improved metabolic health during a dietary intervention in obesity: relationship with gut microbiome richness and ecology. Gut 65(3): 426–436. 10.1136/gutjnl-2014-308778

[10] Everard A, Belzer C, Geurts L, et al. (2013) Cross-talk between Akkermansia muciniphila and intestinal epithelium controls diet-induced obesity. Proc Natl Acad Sci U S A 110(22): 9066–9071. 10.1073/pnas.1219451110

[11] Depommier C, Everard A, Druart C, et al. (2019) Supplementation with Akkermansia muciniphila in overweight and obese human volunteers: a proof-of-concept exploratory study. Nat Med 25(7): 1096–1103. 10.1038/s41591-019-0495-2

[12] Rathod NB, Elabed N, Punia S, Ozogul F, Kim SK, Rocha JM (2023) Recent Developments in Polyphenol Applications on Human Health: A Review with Current Knowledge. Plants (Basel) 12(6). 10.3390/plants12061217

[13] Medina-Larque AS, Rodriguez-Daza MC, Roquim M, et al. (2022) Cranberry polyphenols and agave agavins impact gut immune response and microbiota composition while improving gut barrier function, inflammation, and glucose metabolism in mice fed an obesogenic diet. Front Immunol 13: 871080. 10.3389/fimmu.2022.871080

[14] Barber TM, Kabisch S, Randeva HS, Pfeiffer AFH, Weickert MO (2022) Implications of Resveratrol in Obesity and Insulin Resistance: A State-of-the-Art Review. Nutrients 14(14). 10.3390/nu14142870

[15] Fukizawa S, Yamashita M, Wakabayashi KI, et al. (2020) Anti-obesity effect of a hop-derived prenylflavonoid isoxanthohumol in a high-fat diet-induced obese mouse model. Biosci Microbiota Food Health 39(3): 175–182. 10.12938/bmfh.2019-040

[16] Watanabe S, Tsuneyama K (2012) Cattle bile but not bear bile or pig bile induces lipid profile changes and fatty liver injury in mice: mediation by cholic acid. J Toxicol Sci 37(1): 105–121. 10.2131/jts.37.105

[17] Possemiers S, Heyerick A, Robbens V, De Keukeleire D, Verstraete W (2005) Activation of proestrogens from hops (Humulus lupulus L.) by intestinal microbiota; conversion of isoxanthohumol into 8-prenylnaringenin. J Agric Food Chem 53(16): 6281–6288. 10.1021/jf0509714

[18] Fujisaka S, Avila-Pacheco J, Soto M, et al. (2018) Diet, Genetics, and the Gut Microbiome Drive Dynamic Changes in Plasma Metabolites. Cell Rep 22(11): 3072–3086. 10.1016/j.celrep.2018.02.060

[19] Miyata S, Inoue J, Shimizu M, Sato R (2015) Xanthohumol Improves Diet-induced Obesity and Fatty Liver by Suppressing Sterol Regulatory Element-binding Protein (SREBP) Activation. J Biol Chem 290(33): 20565–20579. 10.1074/jbc.M115.656975

[20] Logan IE, Shulzhenko N, Sharpton TJ, et al. (2021) Xanthohumol Requires the Intestinal Microbiota to Improve Glucose Metabolism in Diet-Induced Obese Mice. Mol Nutr Food Res 65(21): e2100389. 10.1002/mnfr.202100389

[21] Caballero I, Agut M, Armentia A, Blanco CA (2009) Importance of tetrahydroiso alpha-acids to the microbiological stability of beer. J AOAC Int 92(4): 1160–1164

[22] Stevens JF, Taylor AW, Clawson JE, Deinzer ML (1999) Fate of xanthohumol and related prenylflavonoids from hops to beer. J Agric Food Chem 47(6): 2421–2428. 10.1021/jf990101k

[23] Yamashita M, Fukizawa S, Nonaka Y (2020) Hop-derived prenylflavonoid isoxanthohumol suppresses insulin resistance by changing the intestinal microbiota and suppressing chronic inflammation in high fat diet-fed mice. Eur Rev Med Pharmacol Sci 24(3): 1537–1547. 10.26355/eurrev_202002_20212

[24] Fukizawa S, Yamashita M, Fujisaka S, Tobe K, Nonaka Y, Murayama N (2020) Isoxanthohumol, a hop-derived flavonoid, alters the metabolomics profile of mouse feces. Biosci Microbiota Food Health 39(3): 100–108. 10.12938/bmfh.2019-045

[25] Sjostrom L, Rissanen A, Andersen T, et al. (1998) Randomised placebo-controlled trial of orlistat for weight loss and prevention of weight regain in obese patients. European Multicentre Orlistat Study Group. Lancet 352(9123): 167–172. 10.1016/s0140-6736(97)11509-4

[26] Kazmi I, Afzal M, Rahman S, Iqbal M, Imam F, Anwar F (2013) Antiobesity potential of ursolic acid stearoyl glucoside by inhibiting pancreatic lipase. Eur J Pharmacol 709(1-3): 28–36. 10.1016/j.ejphar.2013.02.032

[27] Rodriguez-Daza MC, Pulido-Mateos EC, Lupien-Meilleur J, Guyonnet D, Desjardins Y, Roy D (2021) Polyphenol-Mediated Gut Microbiota Modulation: Toward Prebiotics and Further. Front Nutr 8: 689456. 10.3389/fnut.2021.689456

[28] Plovier H, Everard A, Druart C, et al. (2017) A purified membrane protein from Akkermansia muciniphila or the pasteurized bacterium improves metabolism in obese and diabetic mice. Nat Med 23(1): 107–113. 10.1038/nm.4236

[29] Depommier C, Van Hul M, Everard A, Delzenne NM, De Vos WM, Cani PD (2020) Pasteurized Akkermansia muciniphila increases whole-body energy expenditure and fecal energy excretion in diet-induced obese mice. Gut Microbes 11(5): 1231–1245. 10.1080/19490976.2020.1737307

[30] van der Lugt B, van Beek AA, Aalvink S, et al. (2019) Akkermansia muciniphila ameliorates the age-related decline in colonic mucus thickness and attenuates immune activation in accelerated aging Ercc1 (-/Delta7) mice. Immun Ageing 16: 6. 10.1186/s12979-019-0145-z

[31] Liu MJ, Yang JY, Yan ZH, et al. (2022) Recent findings in Akkermansia muciniphila-regulated metabolism and its role in intestinal diseases. Clin Nutr 41(10): 2333–2344. 10.1016/j.clnu.2022.08.029

[32] Cani PD, Amar J, Iglesias MA, et al. (2007) Metabolic endotoxemia initiates obesity and insulin resistance. Diabetes 56(7): 1761–1772. 10.2337/db06-1491

[33] Pussinen PJ, Havulinna AS, Lehto M, Sundvall J, Salomaa V (2011) Endotoxemia is associated with an increased risk of incident diabetes. Diabetes Care 34(2): 392–397. 10.2337/dc10-1676

[34] Lobo MV, Huerta L, Ruiz-Velasco N, et al. (2001) Localization of the lipid receptors CD36 and CLA-1/SR-BI in the human gastrointestinal tract: towards the identification of receptors mediating the intestinal absorption of dietary lipids. J Histochem Cytochem 49(10): 1253–1260. 10.1177/002215540104901007

[35] Lynes M, Narisawa S, Millan JL, Widmaier EP (2011) Interactions between CD36 and global intestinal alkaline phosphatase in mouse small intestine and effects of high-fat diet. Am J Physiol Regul Integr Comp Physiol 301(6): R1738–1747. 10.1152/ajpregu.00235.2011

[36] Little TJ, Isaacs NJ, Young RL, et al. (2014) Characterization of duodenal expression and localization of fatty acid-sensing receptors in humans: relationships with body mass index. Am J Physiol Gastrointest Liver Physiol 307(10): G958–967. 10.1152/ajpgi.00134.2014

[37] Nassir F, Wilson B, Han X, Gross RW, Abumrad NA (2007) CD36 is important for fatty acid and cholesterol uptake by the proximal but not distal intestine. J Biol Chem 282(27): 19493–19501. 10.1074/jbc.M703330200

[38] Drover VA, Nguyen DV, Bastie CC, et al. (2008) CD36 mediates both cellular uptake of very long chain fatty acids and their intestinal absorption in mice. J Biol Chem 283(19): 13108–13115. 10.1074/jbc.M708086200

[39] Sekar R, Chow BK (2014) Secretin receptor-knockout mice are resistant to high-fat diet-induced obesity and exhibit impaired intestinal lipid absorption. FASEB J 28(8): 3494–3505. 10.1096/fj.13-247536

[40] Demenis C, McLaughlin J, Smith CP (2017) Sulfated Cholecystokinin-8 Promotes CD36-Mediated Fatty Acid Uptake into Primary Mouse Duodenal Enterocytes. Front Physiol 8: 660. 10.3389/fphys.2017.00660

[41] Kuang Z, Wang Y, Li Y, et al. (2019) The intestinal microbiota programs diurnal rhythms in host metabolism through histone deacetylase 3. Science 365(6460): 1428–1434. 10.1126/science.aaw3134

[42] Stojanovic O, Altirriba J, Rigo D, et al. (2021) Dietary excess regulates absorption and surface of gut epithelium through intestinal PPARalpha. Nat Commun 12(1): 7031. 10.1038/s41467-021-27133-7

[43] Watanabe Y, Fujisaka S, Ikeda K, et al. (2021) Gut microbiota, determined by dietary nutrients, drive modification of the plasma lipid profile and insulin resistance. iScience 24(5): 102445. 10.1016/j.isci.2021.102445

[44] Kawano Y, Edwards M, Huang Y, et al. (2022) Microbiota imbalance induced by dietary sugar disrupts immune-mediated protection from metabolic syndrome. Cell 185(19): 3501–3519 e3520. 10.1016/j.cell.2022.08.005

