## Supplementary material for "Isoxanthohumol improves obesity and glucose metabolism via inhibiting intestinal lipid absorption with a bloom of *Akkermansia muciniphila* in mice": Primer

| sybrgreen | Sequence | company |
|---|---|---|
| β-actin | Forward:GCCGGGACCTGACAGACTAC<br>Reverse: AACCGCTCGTTGCCAATAGT | Thermo Fisher Scientific |
| CD11c | Forward:TGTTTGAGTGTCAGGAGCAGG<br>Reverse: GGTCACCTAGTTGGGTCTTG | Thermo Fisher Scientific |
| TNFa | Forward: ACGGCATGGATCTCAAAGAC<br>Reverse: AGATAGCAATCGGCTGACG | Thermo Fisher Scientific |
| GAPDH | Forward:GTCAACGGATTTGGTCGTATTGGG<br>Reverse: TGCCATGGGTGGAATCATATTGG | Thermo Fisher Scientific |
| F4/80 | Forward: CTGGGATCCTACAGCTGCTC<br>Reverse: AGGAGCCTGGTACATTGGTG | Thermo Fisher Scientific |
| UniF340 | ACTCCTACGGGAGGCAGCAGT | Thermo Fisher Scientific |
| UniR514 | ATTACCGCGGCTGCTGGC | Thermo Fisher Scientific |
| CD36 | Forward: ACAGCTGCCTTCTGAAATGTGTGGA<br>Reverse: TTTTCTACGTGGCCCGGTTCTAATTCA | Thermo Fisher Scientific |
| LPL | Forward: TCAGCTGTGTCTTCAGGGGT<br>Reverse: TTTGGCTCCAGAGTTTGACC | Thermo Fisher Scientific |
| NCP1L1 | Forward: CCTAAAGGGGGCCTAGCAGC<br>Reverse: TGCCCGGAGAGCTTCTGTAA | Thermo Fisher Scientific |
| ABCA1 | Forward: GGAAGGGACAAATTGTGCTG<br>Reverse: TGCACAAGGTCCTGAGAATG | Thermo Fisher Scientific |
| ABCG5 | Forward: AGCCATTCACAAACACCTCC<br>Reverse: TCCTTGTACATCGAGAGTGGC | Thermo Fisher Scientific |
| ABCG8 | Forward: TACTTCAGGATGCTTCGCAGG<br>Reverse: TGCTATGAGACCTCCAGGGT | Thermo Fisher Scientific |
| FABP1 | Forward:TGATGTCCTTCCCTTTCTGG<br>Reverse: GCAGAGCCAGGAGAACTTTG | Thermo Fisher Scientific |
| FATP4 | Forward: TTGCAAGTCCCATCAGCAAC<br>Reverse: AACAGCGGCTCTTTCACAAC | Thermo Fisher Scientific |
| Glut5 | Forward:AGCCATCAACAAGGCAGAAC<br>Reverse: CTGCCGTAGAAGACACCACA | Thermo Fisher Scientific |
| PGC1α | Forward: TGATGTGAATGACTTGGATACAGACA<br>Reverse: GCTCATTGTTGTACTGGTTGGATATG | Thermo Fisher Scientific |
| PGC1β | Forward: TCCTGTAAAAGCCCGGAGTAT<br>Reverse: GCTCTGGTAGGGGCAGTGA | Thermo Fisher Scientific |
| PPARα | Forward:TCAGGGTACCACTACGGAGTTCA<br>Reverse: CCGAATAGTTCGCCGAAAGA | Thermo Fisher Scientific |
| ERRα | Forward: CAGCTGTACTCGATGCTCCC<br>Reverse: AGCTCTCTACCCAAACGCCT | Thermo Fisher Scientific |
| A.muciniphila | Forward:CAGCACGTGAAGGTGGGGAC<br>Reverse: CCTTGCGGTTGGCTTCAGAT | Thermo Fisher Scientific |
| SGLT1 | Forward:TCTTCGTCATCAGCGTCATC<br>Reverse: AGGTCGATTCGCTCTTCCTT | Thermo Fisher Scientific |
| PEPT1 | Forward:CGCTTGCCCAAATGTCTC<br>Reverse:CGGTGACCCTGCTCAAAA | Thermo Fisher Scientific |
| Adiponectin | Forward: TGTTCCTCTTAATCCTGCCCA<br>Reverse: CCAACCTGCACAAGTTCCCTT | Thermo Fisher Scientific |
| Glut1 | Forward: AGAACCAATGGCGGCGGTCC<br>Reverse: GCCCGTCACCTTCTTGCTGCT | Thermo Fisher Scientific |
| UCP-1 | Forward: GGCCTCTACGACTCAGTCCA<br>Reverse: TAAGCCGGCTGAGATCTTGT | Thermo Fisher Scientific |
| CideA | Forward: CTAGCACCAAAGGCTGGTTC<br>Reverse: CACGCAGTTCCCACACACTC | Thermo Fisher Scientific |
| LBP | Forward: GGCTCTGCAGAGAGACCTGTACAA<br>Reverse: TAGTTAAGGAATGCCTGGAACAGG | Thermo Fisher Scientific |
| PDK4 | Taqman primer | Thermo Fisher Scientific |
