## Supplementary material for "Isoxanthohumol improves obesity and glucose metabolism via inhibiting intestinal lipid absorption with a bloom of *Akkermansia muciniphila* in mice": Materials

KEY RESOURCES TABLE

REGENT or RESOURCE

| **Chemicals** |
|---|
| Ampicillin |
| Neomycin |
| Metronidazole |
| Vancomycin |
| Brain Heart Infusion |
| Hemin |
| Menadion |
| L-Cysteine |

| **Critical Commercial Assays** |
|---|
| Rneasy Mini Kit (50) |
| 5x Primescript RT Master MIX perfect Realtime |
| PowerFecal DNA kit |
| Triglyceride Colorimetric Assay |
| TB Green Fast qPCR Mix |
| Triglyceride Colorimetric Assay Kit |
| Fecal Mucin Assay Kit |

| **Experimental models** |
|---|
| Mouse C57BL/6J, specific pathogen-free (male) |

| **Oligonnucleotidea** |
|---|
| Primer sequences used for RT-qPCR |

| **Software and Algorithm** |
|---|
| GraphPad Prism 8 |
| FlowJo_v10.6.1 |

| SOURCE | IDENRIER |
|---|---|
| SIGMA-ALDICH | #019M4771V |
| SIGMA-ALDICH | #SLBN5615V |
| SIGMA-ALDICH | #MKCJ4156 |
| LKT LAB | #V0252 |
| BD BBL | 63-6530-20 |
| SIGMA-ALDICH | H9039 |
| SIGMA-ALDICH | M9429 |
| SIGMA-ALDICH | 168149 |

| | |
|---|---|
| QIAGEN | 74104 |
| takara bio | RR036A |
| QIAGEN | 12830-50 |
| Cayman | 10010303 |
| takara bio | RR430A |
| Cayman Chemical Company | 10010303 |
| COSMO BIO CO., LTD. | CSR-FFA-MU-K01 |

| | |
|---|---|
| Japan SLC,Inc. | N/A |

| | |
|---|---|
| See Table | N/A |

| | |
|---|---|
| GraphPad Prism | N/A |
| Becton, Dickinson and Company | N/A |
